# ATHENA: Automatically Tracking Hands Expertly with No Annotations

**DOI:** 10.1101/2025.08.12.669753

**Authors:** Daanish M. Mulla, Mario Costantino, Erez Freud, Jonathan A. Michaels

## Abstract

Studying naturalistic hand behaviours is challenging due to the limitations of conventional marker-based motion capture, which can be costly, time-consuming, and encumber participants. While markerless pose estimation exists – an accurate, off-the-shelf solution validated for hand-object manipulation is needed. We present ATHENA (Automatically Tracking Hands Expertly with No Annotations), an open-source, Python-based toolbox for 3D markerless hand tracking. To validate ATHENA, we concurrently recorded hand kinematics using ATHENA and an industry-standard optoelectronic marker-based system (OptiTrack). Participants performed unimanual, bimanual, and naturalistic object manipulation and we compared common kinematic variables like grip aperture, wrist velocity, index metacarpophalangeal flexion, and bimanual span. Our results demonstrated high spatiotemporal agreement between ATHENA and OptiTrack. This was evidenced by extremely high matches (R^2^ > 0.90 across the majority of tasks) and low root mean square differences (< 1 cm for grip aperture, < 4 cm/s for wrist velocity, and < 5-10° for index metacarpophalangeal flexion). ATHENA reliably preserved trial-to-trial variability in kinematics, offering identical scientific conclusions to marker-based approaches, but with significantly reduced financial and time costs and no participant encumbrance. In conclusion, ATHENA is an accurate, automated, and easy-to-use platform for 3D markerless hand tracking that enables more ecologically valid motor control and learning studies of naturalistic hand behaviours, enhancing our understanding of human dexterity.

## Introduction

The human hand allows for dexterous manipulation unparalleled by any other living organism. As a result, humans can perform a rich repertoire of tasks using their hands, ranging from activities involving delicate finger control to forceful grasps. Despite the importance of hands in our evolutionary history and every facet of our lives today, from nonverbal communication to activities of daily living, surprisingly little is known about how humans acquire manipulative skills (Krakauer et al., 2019; Longo, 2025; Xu et al., 2024). Much of the motor learning literature on the upper limb is focused on reaching, with what little there is known about dexterous hand abilities limited to free-hand finger movements and grasps (Sobinov & Bensmaia, 2021; Valero-Cuevas & Santello, 2017). To understand neural control of the hand, we need to study naturalistic hand behaviours that reflect how individuals learn to perform complex skills in real-world settings and how these movements may be altered across neurodevelopmental or neurodegenerative disorders (Freud et al., 2025).

The primary challenge in studying naturalistic hand behaviours is recording hand movements (Valero-Cuevas & Santello, 2017). Conventionally, hand movements are recorded through optoelectronic marker-based motion capture systems, manual annotation of video recordings, or instrumented gloves (Keenan et al., 2009; Santello et al., 1998; van Beek et al., 2018; Yan et al., 2020, 2022; Yao et al., 2021). These systems have several limitations including financial costs, markers / sensors encumbering participants, and the collection and processing of data is time-consuming. These limitations have restricted researchers to studying hand behaviours during simple movements (e.g., individuated finger movement, grasps) within constrained in-lab settings that fail to capture the rich breadth of hand capabilities such as observed during complex object manipulation. Recent progress in computer vision has allowed researchers to start to overcome many of these limitations by training machine learning models to automatically identify hand key points from video recordings and enable hand pose estimation (i.e., markerless tracking). For example, markerless tracking was combined with high-density electromyographic recordings to study recruitment of motor unit populations during single and multi-digit finger movements (Oßwald et al., 2025). Although toolboxes are available for training models to perform markerless pose estimation (Mathis et al., 2018), a barrier for implementation is the initial requirement for manually annotating training images. Also, pose estimation tools do not always support 3D tracking with multiple cameras. One solution is to combine pre-trained models with a custom or open-source triangulation pipeline, as demonstrated for free-hand single- and multi-finger motion (Firouzabadi et al., 2024; Mulla et al., 2025). However, the feasibility of markerless methods for hand pose estimation during object manipulation and their tracking accuracy, as validated against the “industry standard” optoelectronic marker-based methods, have not been assessed. Hence, there is a need for an accurate, validated, off-the-shelf solution for an end-to-end markerless 3D tracking of the hands using multiple video camera recordings.

The aim of our work was to share a user-friendly, accurate solution for markerless tracking of the hands. Here, we present ATHENA (Automatically Tracking Hands Expertly with No Annotations), an open-source Python based toolbox for performing 3D markerless tracking of the hands. In this paper, we describe the development and experimental validation of our toolbox. To validate ATHENA, we compared the accuracy of hand kinematics from a markerless system against an optoelectronic marker-based system during simple unimanual and bimanual reach-and grasp tasks, as well as complex naturalistic behaviours with hand-object manipulation representative of activities of daily living.

## Materials and Methods

### Description of ATHENA

An overview of ATHENA’s workflow is shown in Figure 1. The required inputs are synchronized video recordings along with the intrinsic and extrinsic parameters of each camera. ATHENA uses MediaPipe Solutions (Lugaresi et al., 2019; Zhang et al., 2020) to identify 2D locations of 42 key points bimanually across the hands (*Hand Landmarker model*) and 33 key points across the entire body (*Pose Landmarker model*) for each frame across all video recordings. MediaPipe was chosen as it has a pre-trained model for tracking hand key points compared to other open-source toolboxes where users are required to train their own models (e.g., DeepLabCut). Users are provided options for setting the minimum hand and pose detection and tracking confidences within MediaPipe. For the results presented in this paper, all confidence thresholds were set to 0.90. ATHENA supports parallel processing, with multiple cameras processed in parallel across different CPU cores. The intrinsic camera parameters are used at this stage for undistorting the images. MediaPipe’s hand model can be susceptible to handedness errors (right versus left hand). To identify and correct for handedness errors, the proximity of the wrist landmarks of the hand model for each camera view is evaluated against (1) the wrist landmarks of the pose model to ensure alignment in handedness between models, and (2) the reprojected 2D wrist location following triangulation for circumstances where the pose model did not detect the arms. Following correction of handedness, a direct linear transform is applied to all 2D key point locations to triangulate to world 3D coordinates using singular value decomposition. The final 3D positions are smoothed using a Savitzky-Golay filter, with the cut-off frequency choice given to the user. For the findings in the current paper, a 10 Hz cut-off frequency was set unless otherwise specified. ATHENA requires minimal dependencies, with optional GPU usage and no required machine learning libraries. Upon installation, users are presented with a graphical user interface allowing them to select required inputs (camera calibrations, video recordings) and optional processing choices (MediaPipe detection and tracking confidences, filter cut-off). Installation instructions and documentation are available on GitHub at https://github.com/neural-control-and-computation-lab/athena.

**Figure 1.**
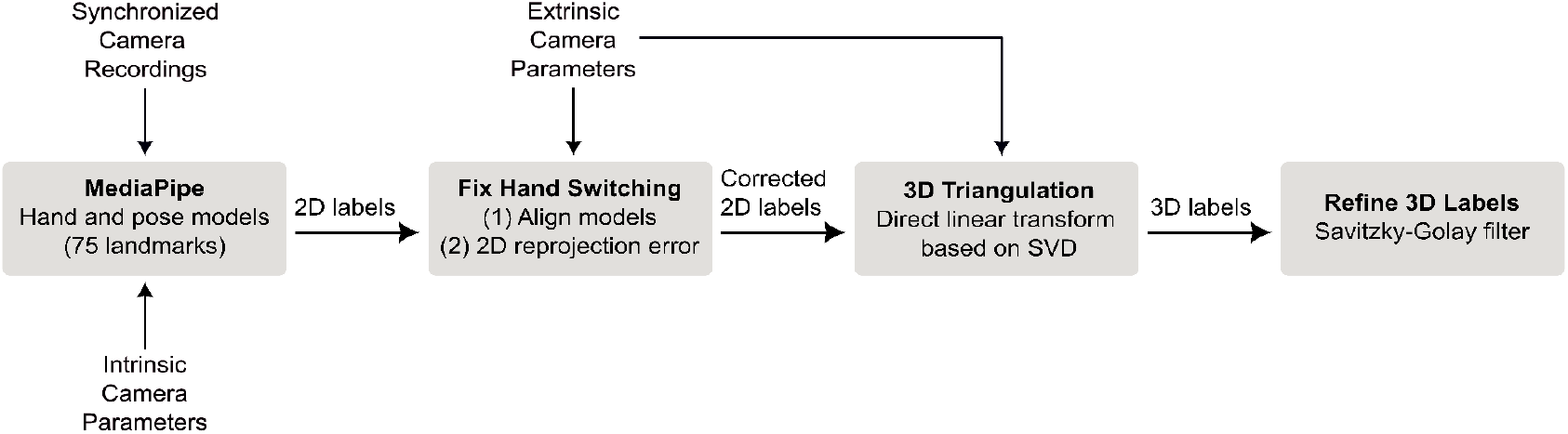
A schematic of ATHENA’s workflow. Inputs required include synchronized camera recordings and camera intrinsic and extrinsic parameters.

### Kinematic Recordings

To validate ATHENA, we conducted an experiment where hand movements were concurrently recorded using a markerless and marker-based optoelectronic motion capture system (Figure 2a). The cameras from both systems were mounted on a 2 × 2 × 2 m frame. The markerless system comprised 8 video cameras (BFS-U3-13Y3C-C, Teledyne FLIR, USA) with lenses (Kowa LM5JC1M) recorded at 1280 × 1024 resolution and sampled at 60 Hz with an exposure time of 8 ms. All data were recorded and processed on a PC with an AMD Ryzen 9 7950×3D processor and an NVIDIA RTX 3060 graphics card. Video recordings and camera calibration were performed using JARVIS Mocap (https://jarvis-mocap.github.io/jarvis-docs/). Reprojection error across calibration for all cameras was less than 0.5 pixels. Video recordings were processed using ATHENA to obtain 3D markerless kinematics. Marker-based kinematics were collected using a 5-camera passive motion capture system sampled at 100 Hz (Prime 13w, Natural Point DBA OptiTrack, USA). For unimanual reach and grasp tasks (see below), we used a 4 and 9 reflective marker set, with all markers affixed to the dorsal aspect of the right hand (Figure 2a). The 4-marker set included: (1) wrist – midway between the radial and ulnar styloid processes, (2) thumb trapeziometacarpal joint, (3) thumb fingertip, and (4) index fingertips. In addition to these markers, the 9-marker set included: (5) thumb metacarpophalangeal joint, (6) thumb interphalangeal joint, (7) index metacarpophalangeal joint, (8) index proximal interphalangeal joint, and (9) index distal interphalangeal joint. For bimanual reach and grasp tasks, the 4-marker set was repeated across both hands for a total of 8 reflective markers (Figure 3a). For naturalistic behaviours, we used the 4-marker set on the right hand (Figure 4a). All reflective markers were 3 mm hemispheres.

**Figure 2.**
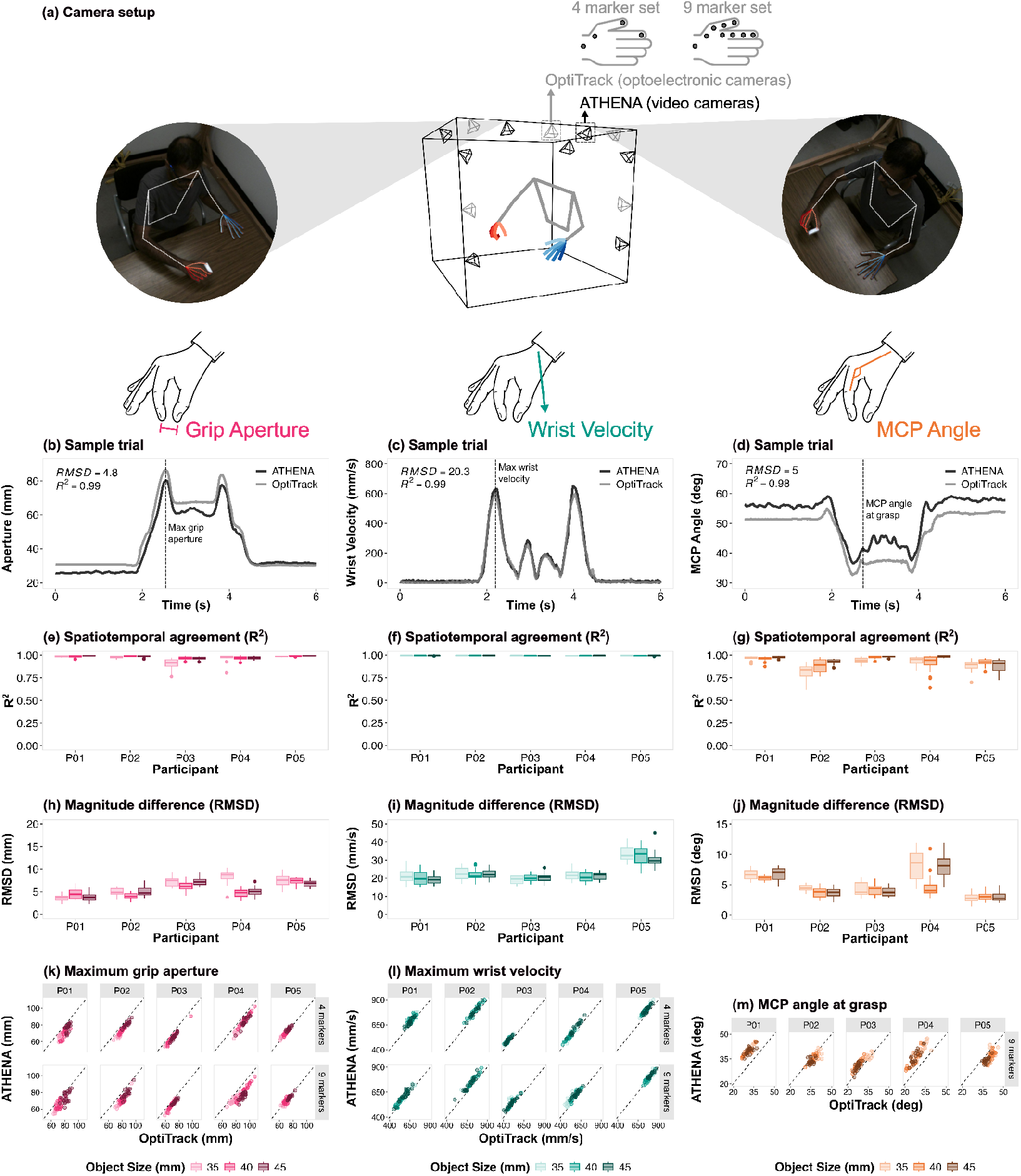
Comparisons between ATHENA and OptiTrack during unimanual reach and grasp trials. (a) Visual depiction of the experimental setup with concurrent recordings from 8 video cameras (ATHENA) and 5 optoelectronic cameras (OptiTrack). OptiTrack recordings for bimanual tasks were made with a 4 and 9 marker set. Still frames from video camera recordings are shown with ATHENA’s predictions overlaid. (b-d) Signal traces from a sample trial comparing (b) grip aperture, (c) wrist velocity, and (d) MCP angle between ATHENA (black) and OptiTrack (grey), with RMSD and R^2^ values for the trial reported on the top left corner. The distribution of (e-g) R^2^ and (h-j) RMSD values across all trials. Scatter plots for (k) grip aperture, (l) wrist velocity, and (m) MCP angle between ATHENA and OptiTrack across all trials separated by participants (P01-P05) and number of markers (4 and 9). The dotted line is the line of equality. The colour shades in panels (e-m) represent different object sizes.

**Figure 3.**
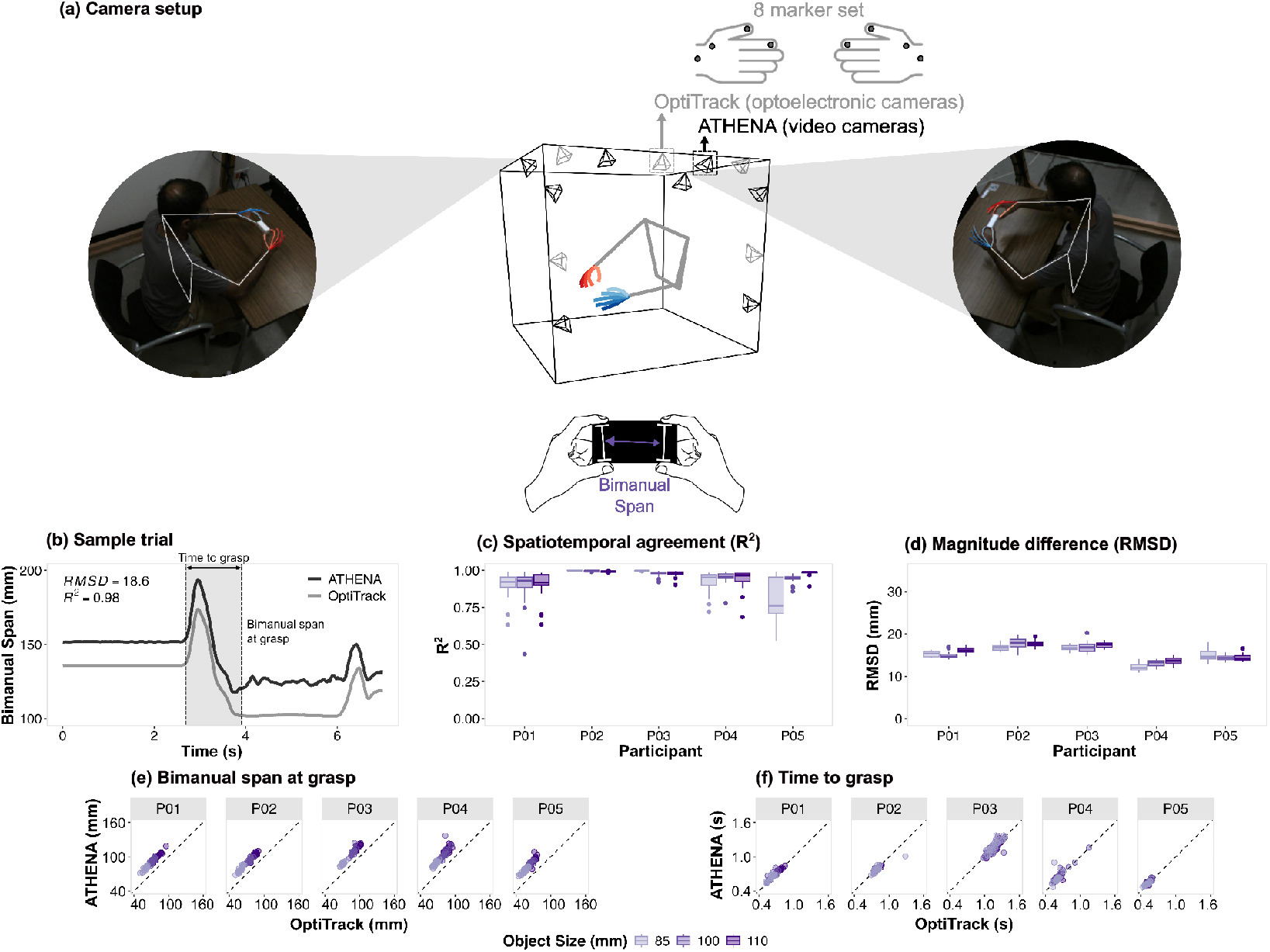
Comparisons between ATHENA and OptiTrack during bimanual reach and grasp trials. (a) Visual depiction of the experimental setup. OptiTrack recordings for bimanual tasks were made with an 8 marker set (4 markers on each hand). Still frames from video camera recordings are shown with ATHENA’s predictions overlaid. (b) Signal traces from a sample trial between ATHENA (black) and OptiTrack (grey), with RMSD and R^2^ values for the trial reported on the top left corner. The distribution of (c) R^2^ and (d) RMSD values across all trials. Scatterplots for (e) bimanual span at grasp and (f) time to grasp from the start of the reach between ATHENA and OptiTrack across all trials separated by participants (P01-P05). The dotted line is the line of equality. The shades of purple in panels (c-f) represent different object sizes.

**Figure 4.**
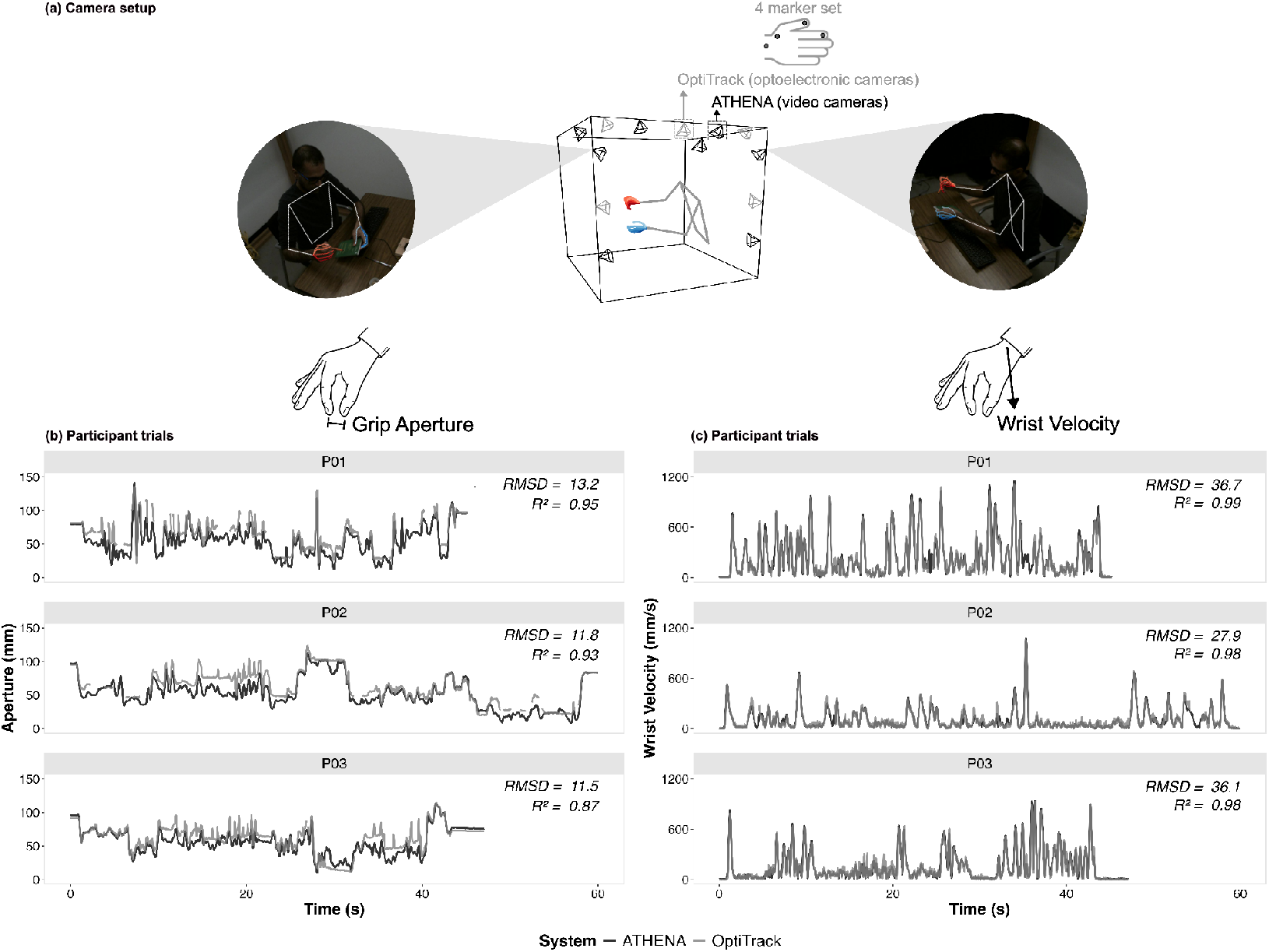
Comparisons between ATHENA and OptiTrack during naturalistic tasks. (a) Visual depiction of the experimental setup. OptiTrack recordings for naturalistic tasks were made with a 4 marker set. Still frames with ATHENA’s predictions overlaid are shown while the participant was grasping a book (left hand) and pen (right hand). (b) Grip aperture and (c) wrist velocity comparisons between ATHENA (black) and OptiTrack (grey) for three participants (rows: P01-P03). Data is plotted across the entire trial (40-60 second durations), with RMSD and R^2^ between the systems for each trial reported on the top right corner of each graph.

### Experimental Procedures

Participants gave verbal and written consent and performed three sets of experimental tasks: (1) unimanual reach and grasp (N=2M/3F; age=23-35 years old), (2) bimanual reach and grasp (N=2M/3F; age=23-35 years old), and (3) naturalistic behaviour (N=3M; age=21-35 years old). The experimental protocol was approved by the Human Participants Review Committee (HRPC) at York University. Participants had no history of neurological conditions or upper extremity musculoskeletal injuries. For unimanual reach and grasp tasks, participants were instructed to use their right hand to grasp a target object with a two-finger pinch grip across the length of the object and lift it above the table before setting it down in the same position (Freud et al., 2025). Participants started and ended each trial at a “home” position at the edge of the table. The target object was a rectangular prism with the wide side facing away from the participant and varying in size (lengths of 35, 40, 45 mm). Twenty trials were recorded for each size and all trials were repeated across three marker set conditions: without any markers and with both the 4- and 9-marker sets (total of 180 trials per participant). The bimanual reach and grasp tasks were conducted similarly; however, the wide side of the target object (lengths of 85, 100, 110 m) faced the participant. Participants were instructed to use two hands to pinch grip each end of the object and lift it above the table. The bimanual reach and grasp tasks were conducted twice: without any markers and with the 8-marker set (total of 120 trials per participant). The order of the unimanual and bimanual reach and grasp tasks were block randomized, such that all trials for a given marker set and task combination (0-, 4-, 9-marker unimanual and 0-, 8-marker bimanual) were completed before proceeding to the next combination. Within each marker set and task combination, the order of the object sizes were block randomized such that all trials for a given object size were completed prior to moving to the next object size. For the naturalistic behaviours, a set of daily-use objects were placed on a table (e.g., keyboard, light bulb fixture, screwdriver, pen, book, mug). Participants were instructed to freely manipulate the objects as they wished during a 40-60 second trial. A single trial was recorded per participant.

### Data Analysis

Comparisons between the markerless (i.e., ATHENA) and marker-based (i.e., OptiTrack) systems were based on 4 kinematic variables: (1) grip aperture, (2) wrist linear velocity, (3) index metacarpophalangeal (MCP) flexion, and (4) bimanual span. These variables were chosen to encompass the range of interest (distances, higher order kinematic derivatives, joint angles) among movement scientists. Grip aperture was defined as the distance between the thumb and index fingertips. Index MCP flexion was determined using the dot product of the vectors comprising the index metacarpal and proximal phalanx (Mulla et al., 2025) and was only quantified for the 9-marker set unimanual reach and grasp trials. Bimanual span was defined as the distance between the midpoint of the thumb and index fingertips across the two hands and was only quantified for the bimanual reach and grasp trials. To match the sampling rates across systems, the markerless kinematics were upsampled to 100 Hz. As the markerless and marker-based systems were not hardwired to record synchronously, a cross-correlation was performed on the grip aperture to identify the time shift and post-hoc synchronize the systems. To determine spatiotemporal agreement between the markerless and marker-based systems, the root mean square difference (RMSD) and correlation coefficient (R^2^) for all kinematic variables were calculated for each trial. As discrete metrics (e.g., peaks, instantaneous timing-related information) rather than continuous 1D signals may be of interest to some researchers (Aglioti et al., 1995; Freud et al., 2025; Marras & Schoenmarxlin, 1993; Mulla et al., 2025), we also extracted 4 kinematic features during the reaching phase of the unimanual and bimanual reach and grasp movements: (1) maximum grip aperture, (2) maximum wrist velocity, (3) index MCP flexion at grasp, and (4) bimanual span at grasp. The start of the reaching movement was defined as the frame following twenty consecutive frames where the wrist velocity exceeded 20 mm/s. The end of the reaching movement (i.e., start of the grasping movement) was defined as the frame where the wrist velocity reached a minimum between the first (reaching) and second (lifting) local maximums in the wrist velocity. A total of 20 markered unimanual and bimanual reach and grasp trials were discarded due to marker dropout and movement start criteria was not met in one or both of the systems. Unless otherwise specified, the quantitative comparisons between the systems for the unimanual reach and grasp tasks are reported for the 4-marker set for grip aperture and wrist velocity, and the 9-marker set for index MCP flexion.

## Results

Comparisons between the markerless (i.e., ATHENA) and marker-based (i.e., OptiTrack) systems are shown in Figures 2-4 for the unimanual reach and grasp, bimanual reach and grasp, and naturalistic behaviours, respectively. For all tasks – unimanual reach and grasp (Figure 2b-d), bimanual reach and grasp (Figure 3b), and naturalistic behaviour (Figure 4b,c) – we observed qualitative similarities in the 1D kinematic signals between the markerless and marker-based kinematics.

Visual inspections were supported by formal quantitative comparisons of the continuous spatiotemporal data, with high R^2^ and low RMSD values observed for all kinematic variables during the simpler unimanual and bimanual reach and grasp tasks. During the unimanual reach and grasp, closest agreement was observed for the wrist velocity (median [interquartile range] R^2^ = 0.99 [0.99-1.00], RMSD = 21.5 mm/s [19.4-25.8]; Figure 2f,i), followed by grip aperture (R^2^ = 0.98 [0.97-0.99], RMSD = 5.8 mm [4.4-7.2]; Figure 2e,h), and then MCP flexion (R^2^ = 0.96 [0.90-0.98], RMSD = 4.4° [3.4-6.1]; Figure 2g,j). The agreement between the systems slightly decreased when increasing the number of markers from 4 to 9 for wrist velocity (RMSD = 24.2 mm/s [21.2-29.6]) and grip aperture (RMSD = 6.6 mm [5.0-8.7]). We observed a strong linear relationship between the two systems for all discrete kinematic metrics – maximum grip aperture (Figure 2k), maximum wrist velocity (Figure 2l), and index MCP flexion at grasp (Figure 2m). During the bimanual reach and grasp, the median [interquartile] R^2^ in bimanual span was 0.98 [0.94-0.99], with RMSD values of 15.5 mm [14.0-16.9] (Figures 3c,d). A strong linear relationship was observed between the two systems in the bimanual span at grasp (Figure 3e) and time to grasp from start to reach (Figure 3f).

Similar close agreement between the systems were observed during naturalistic behaviours. The R^2^ in grip aperture and wrist velocity ranged from 0.87-0.95 and 0.98-0.99, respectively, with RMSD values between 11.5-13.2 mm (grip aperture) and 27.9-36.7 mm/s (wrist velocity) across the participants (Figure 4b,c). Notably, we observed several instances of marker dropout within the marker-based system, particularly for the thumb and index fingertips (see discontinuous grey lines in Figure 4b) due to marker occlusion by objects or other parts of the body. Although occlusion can affect detection and tracking of key points in the markerless system, estimates of key point locations can frequently still be made (tuned by the input confidence thresholds) by the MediaPipe models based on similar postures encountered during the training stage (see continuous black lines in Figure 4b). Further, the lower expense of the markerless systems facilitates the purchase of more cameras, which may help circumvent occlusion issues by offering additional views of the hands.Sensitivity of the spatiotemporal agreement in the 1D kinematic data between the two systems (i.e., R^2^) to the choice of filter cut-off is depicted in Figure 5 for the unimanual and bimanual reach and grasp tasks. Across both tasks and for all kinematic variables, a cut-off frequency less than 5 Hz (i.e., 1 Hz) drastically reduced the correlations between the systems as much of the spatiotemporal movement-related information was attenuated. The correlations were largely invariant to cut-off frequencies over the range of 5-20 Hz, but were sensitive when exceeding 20 Hz for the wrist velocity as was expected because higher-order derivatives are more sensitive to signal noise upon differentiation.

**Figure 5.**
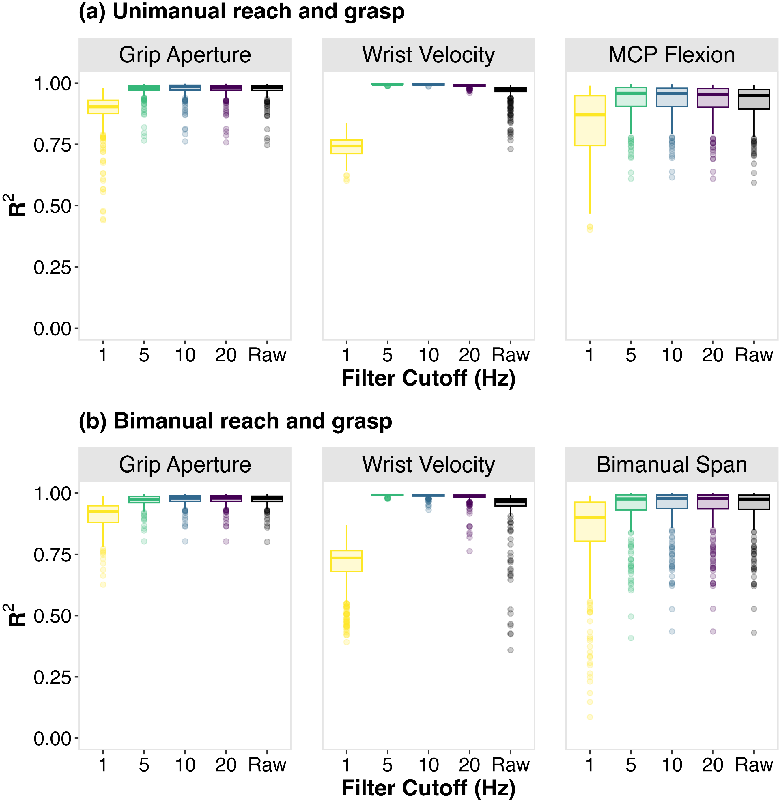
Correlations between ATHENA and OptiTrack for all (a) unimanual and (b) bimanual reach and grasp trials across different Savitzky-Golay filter cutoff frequencies (1–20 Hz or unfiltered [raw]).

## Discussion

In this work, we present ATHENA, a Python-based toolbox for markerless tracking of the hands. We leveraged open source 2D tracking solutions with a simple processing pipeline consisting of correcting hand switching, triangulation, and smoothing to automatically track 3D locations of several key points across the hands and body. As a test of ATHENA’s tracking accuracy, we concurrently recorded with a markerless and industry standard marker-based motion capture system and found comparable results between two systems. Overall, ATHENA provides the scientific community with an accurate solution for obtaining 3D hand kinematic data from raw videos while requiring minimal user input, thereby facilitating studies on the neural control of hand movements.

ATHENA is a general-purpose solution to overcome persistent challenges in tracking 3D hand kinematics. The common use of marker-based motion capture and instrumented gloves for recording hand behaviours are restricted by two main limitations: (1) costs (financial, time), and (2) encumbering participant movements. The use of markers and sensors, particularly for the hands, can be arduous (e.g., up to 30-60 minutes of setup time), error-prone (e.g., soft-tissue motion artefact, identifying landmarks), require time-consuming digitizing steps, and restricts finger motion, thereby affecting the types of movements recorded and the ecological validity of the behaviours captured (Cocchiarella et al., 2016). Further, markers can be uncomfortable for participants with tactile hypersensitivity and lead to study exclusion. ATHENA resolves these challenges by using synchronized video recordings requiring no markers / sensors and no manual processing steps to automatically track hand kinematics. For our markerless motion capture system (8-cameras, 1280×1024 resolution, 60 Hz), ATHENA ran offline at approximately 0.25x real-time (i.e., a one-minute video is fully processed in four minutes, including saving images and videos). Notably, the collection setup used in our validation study are not the minimum requirements. In an earlier version of the toolbox, we demonstrated that a 2-camera system recorded at a 480×640 resolution and sampled at 30 Hz can suffice for tracking hand kinematics during single finger flexion-extension (Mulla et al., 2025). On the other hand, with more unconstrained movements as observed during real-world settings, more cameras may be necessary (Bala et al., 2020). ATHENA was developed with these diverse experimental conditions in mind, providing flexibility to end users in their markerless motion capture system setups (e.g., number of cameras, video resolution, sampling rate) for their own applications.

Across both simple reach and grasp tasks and more complex, naturalistic behaviours, we found relatively small kinematic differences between ATHENA and OptiTrack. There was a high degree of spatiotemporal agreement between systems with R^2^ across most trials exceeding 0.9 and relatively low RMSD values (< 1 cm for grip aperture, < 4 cm/s for wrist velocity, and < 5-10° for index MCP flexion). We observed some systematic differences, primarily for grip aperture with ATHENA estimating apertures consistently less than those tracked by OptiTrack, which can be explained by the location difference of ATHENA’s key point estimation (joint level) versus marker placement (surface of the skin). Nevertheless, strong linear relationships existed across all discrete kinematic metrics (e.g., maximum grip aperture, maximum wrist velocity) between the systems. Thus, despite any small differences in the absolute value of the kinematic metrics between systems, ATHENA can reliably track and preserve relative differences in magnitude between trials (i.e., across repetitions, experimental conditions). This suggests that ATHENA would yield similar study conclusions to conventional marker-based motion capture systems but notably, with a fraction of the financial and time costs and without encumbering participants due to marker placements. Our experimental findings are in line with the similar validation results of markerless motion capture systems in recent years; however, much of these works have focused on lower body and proximal upper limb kinematics (Hansen et al., 2024; Kanko et al., 2021; Uhlrich et al., 2024). Advances in 3D markerless technologies of the hand and fingers have received less attention in the movement sciences, with the few developments to date focused exclusively on free-hand motion and commonly missing validation analyses (Firouzabadi et al., 2024; Gionfrida et al., 2022; Mulla et al., 2024; Oßwald et al., 2025). By enabling users to track hands during object manipulation with kinematic accuracy comparable to industry standard optoelectronic marker-based systems, ATHENA can help researchers overcome the long-standing challenge of recording naturalistic hand behaviours.

A few limitations need to be considered for users interested in ATHENA. The toolbox is designed for human subjects and cannot track non-human primates (e.g., see Bala et al., 2020 and JARVIS mocap for available tracking solutions with pre-trained models for non-human primates). Further, MediaPipe has known challenges for detecting hand landmarks in the presence of gloves or extensive hand tattoos and henna, limiting ATHENA’s generalizability across experimental conditions and participants. The issues of tracking non-human primates and estimating postures for gloved or inked / dyed hands may be resolved in the future through re-training MediaPipe models. Concurrent validation was only performed for hand kinematics. Although ATHENA outputs whole body posture using MediaPipe’s pose model, marker-based motion capture was only performed for the hands and thus, validity of 3D kinematics for the entire body is unknown. Finally, as with any markerless motion capture system, the use of video recordings for kinematic tracking presents important data security and privacy issues that must be proactively considered by users for safe adoption (Armitano-Lago et al., 2022).

In conclusion, we share an easy-to-use and accurate open-source toolbox (ATHENA) for performing automated 3D markerless tracking of the hand using synchronized video recordings. We found close agreement between ATHENA’s kinematics with an industry standard optoelectronic marker-based motion capture system across a series of simple and complex object manipulation tasks. ATHENA is well-suited to facilitate future motor control and learning studies for investigating naturalistic hand behaviours.

## Acknowledgements

We would like to thank Timo Hüser and Benjamin Dann for help with data acquisition software, and Ali Ghavampour for work on an earlier version of our hardware setup.

## Disclaimers

The content is solely the responsibility of the authors and does not necessarily represent the official views of the institution.

## Author Contributions

- Conceived and designed research: DMM, EF, JAM
- Analyzed data: DMM, JAM
- Performed experiments: DMM, MC
- Interpreted results of experiments: DMM, JAM
- Prepared figures: DMM
- Drafted manuscript: DMM
- Edited and revised the manuscript: DMM, MC, EF, JAM
- Approved final version of the manuscript: DMM, MC, EF, JAM

## Data Availability

Data to reproduce the findings are available upon request.

